# Drug-induced resistance in micrometastases: analysis of spatio-temporal cell lineages

**DOI:** 10.1101/2020.03.01.972406

**Authors:** Judith Pérez-Velázquez, Katarzyna A. Rejniak

**Affiliations:** Mathematical Modeling of Biological Systems, Centre for Mathematical Science, Technical University of Munich, Garching, Germany; H. Lee Moffitt Cancer Center & Research Institute, Integrated Mathematical Oncology Department, Tampa, Florida, USA; University of South Florida, Morsani College of Medicine, Department of Oncologic Sciences, Florida, USA

## Abstract

Resistance to anti-cancer drugs is a major cause of treatment failure. While several intracellular mechanisms of resistance have been postulated, the role of extrinsic factors in the development of resistance in individual tumor cells is still not fully understood. Here we used a hybrid agent-based model to investigate how sensitive tumor cells develop drug resistance in the heterogeneous tumor microenvironment. We characterized the spatio-temporal evolution of lineages of the resistant cells and examined how resistance at the single-cell level contributes to the overall tumor resistance. We also developed new methods to track tumor cell adaptation, to trace cell viability trajectories and to examine the three-dimensional spatio-temporal lineage trees. Our findings indicate that drug-induced resistance can result from cells adaptation to the changes in drug distribution. Two modes of cell adaptation were identified that coincide with microenvironmental niches—areas sheltered by cell micro-communities (protectorates) or regions with limited drug penetration (refuga or sanctuaries). We also recognized that certain cells gave rise to lineages of resistant cells (precursors of resistance) and pinpointed three temporal periods and spatial locations at which such cells emerged. This supports the hypothesis that tumor micrometastases do not need to harbor cell populations with pre-existing resistance, but that individual tumor cells can adapt and develop resistance induced by the drug during the treatment.

## Introduction

Drug resistance is one of the main impediments in effective anti-cancer therapy. While tumors may first respond well to chemotherapeutic agents, they often start growing back during or after the treatment period and become tolerant to the treatment. Several different intrinsic mechanisms of drug resistance have been postulated (1–3), including alteration of drug targets, changes in the expression of efflux pumps, increased ability to repair DNA damage, reduced apoptosis, elevated cell death inhibition, and altered proliferation. Some extrinsic factors contributing to drug resistance have also been postulated. A pivotal role can be played by the tumor microenvironment (4, 5) due to reciprocal communication between tumor cells and the surrounding stromal components. This includes interactions with fibroblasts and emergence of cancer associated fibroblasts, cross-talk with immune cells and sensing cues from the extracellular matrix (ECM), as well as ECM remodeling. Tumor cells can modify their microenvironment by creating specific niches, including pre-cancerous, pre-metastatic or stem cell niches (6–8). Additionally, the changes in tumor vasculature and interstitial fluid pressure may lead to creation of regions that are poorly penetrated by therapeutics, forming drug-limited pharmacologic sanctuaries or refugia (9, 10) that influence tumor response to therapeutics. However these extrinsic factors are still not well understood.

Of particular interest is the heterogeneous and dynamically changing tumor microenvironment. As a result, spatially and temporarily variable gradients of drugs can be formed in the stroma, and tumor cells can be, therefore, exposed to different drug levels during the treatment period. It has been shown experimentally by Wu and colleagues (11) that aggressive breast tumor cells can respond to drug gradients by migrating toward the regions of high concentration of doxorubicin and low cell population, and are able to adapt to high drug levels. Subsequently, they become tolerant to the drug and repopulate the region despite the high drug concentration. Fu and colleagues (12) used mathematical modeling to investigate how the heterogeneity in drug penetration through the microenvironment effects tumor response to treatment. They showed that resistance arises first in cells located in regions with poor drug penetration, named pharmacological sanctuaries, and then populate areas with higher drug levels. Our own research showed that the non-homogeneous drug distribution within the tumor tissue that results in emergence of tissue regions with poor drug penetration but with normal oxygenation levels may lead to the emergence of acquired resistance (13, 14). Similar results were previously generated using different mathematical models. Chisholm et al. (15) investigated transient emergence of a drug tolerant population of cells using models of reversible phenotypic evolution, and concluded that a combination of non-genetic instability, stress-induced adaptation and selection are responsible for the emergence of weakly-proliferative and drug-tolerant tumor cells. Cho and Levy (16) used a continuous model to show that cancer cells of different resistance levels can coexist in spatially-different areas in tumor tissue. Feizabadi (17) used mathematical modeling to show that certain chemotherapy strategies are highly unsuccessful, and even damaging to the patient, under the assumption that the drug can induce resistance during the treatment period. Greene et al. (18, 19) developed mathematical approach to differentiate between spontaneous and induced resistance to drugs and proposed in vitro experiments that can determine weather treatment can induce resistance. The authors also designed optimized treatment protocols that can prolong the time before resistance develops.

Several experimental studies considered scenarios in which resistance is acquired by the tumor cells as a result of their exposure to the drug, either through epigenetic alteration, drug-induced genetic changes or non-genetic phenotype switching. Pisco and colleagues (20, 21) used a combination of laboratory experiments and mathematical modeling to show that the emergence of multi-drug resistance in leukemic cells can be induced by the lasting stress response to the drug. In this case, the tumor cells exploited their phenotypic plasticity by modifying efflux capacity in a non-genetic but inheritable way. Goldman and colleagues (22, 23) showed that exposure of breast tumor cells to high concentration of taxanes can induce phenotypic transitions towards chemotherapy-tolerant stem-like state that can confer drug resistance. Moreover, the authors demonstrated that this adaptive resistance process can be halted by carefully designed order of administered drug combinations. Other examples of drug-induced resistance pointed to modifications in chromatin configuration in lung cancer cells (24, 25), changes in expression of stress adaptation-related proteins in prostate cancer cells (26), or switching to mesenchymal phenotype in melanoma cells (27) as the mechanisms of increased cell tolerance to the drug. In all these studies, the exposure of tumor cells to chemotherapy caused non-genetic changes that allowed the tumor cells to tolerate drug treatment and evade drug-induced death.

In this paper, we used mathematical modeling to examine how individual tumor cells can adapt to alterations in drug distribution within the tumor microenvironment in order to acquire resistance to the drug. By tracking cells individually and reconstructing their behavioral history, we were able to provide insights into the complex spatiotemporal changes that occur in cell microcommunities and to explain how they avoid drug-induced death leading to therapy failure. In particular, we developed a concept of 3D spatio-temporal lineage trees that trace both genealogy and spatial locations of cells that survived the simulated treatment. This is an extension of classical lineage trees used to depict tumor clonal expansion in a form of a flat graph with an initiating cell connected to its children cells, that are connected to their descendants until the terminal nodes are reached (28, 29). The 3D spatio-temporal lineage trees allow us to identify the cells that drive a resistant phenotype in the sense that all their successors are resistant to the drug. The existence of such “special” cells has been reported previously under various names: drivers (30, 31), superstars (32, 33), or starter cells (34). We refer to these cells as precursors of drug resistance. The current study focuses on analyzing the behavior of individual resistant cells which is an extension of our previous work at the population level. This approach allowed us to develop novel evaluation methods, such as the 3D lineage trees, and also to identify the third microenvironmental niche prone to the emergence of resistant cells. Overall, this paper contributes to a better understanding of drug-induced resistance.

## Materials and methods

We used a hybrid multi-cell lattice-free model (*MultiCell-LF*) that combines the off-lattice individual tumor cells with the continuous description of oxygen and a cytotoxic drug. The cells can physically interact with one another, and respond to levels of oxygen and cytotoxic drug absorbed from cell’s vicinity. Low levels of oxygen (hypoxia) result in cell quiescence (35). Exposure to the drug leads to cell damage—while we model this as a generic process, one can consider a more specific processes, such as the DNA damage (genotoxicity (36)) or cell membrane damage (lysis (37)). Moreover, cells can become more tolerant to the drug they are exposed to, as shown in (21, 25). This cell’s response to different levels of oxygen and drug is a mechanism of cell adaptation to the microenvironment.

### Drug and oxygen kinetics

The model is defined on a small patch of the tumor tissue with four irregularly positioned stationary vessels (Figure 1AB). Both drug γ and oxygen *ξ* are intravenously supplied, diffuse through the tissue, are absorbed by the cells, and the drug is subject to decay. We model a small drug molecule of diffusivity comparable to oxygen diffusion (38) and with the same supply rate from the vessels. However, the drug is absorbed by the cells twice faster than oxygen (38). Drug *γ* and oxygen *ξ* kinetics are given by the following equations:

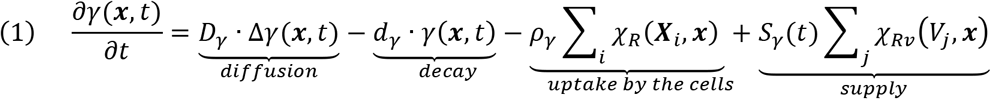

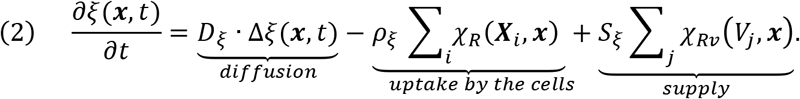

where, *D*_*γ*_ and *D*_*ξ*_ are the drug and oxygen diffusion coefficients, *ρ*_*γ*_ and *ρ*_*ξ*_ are the drug and oxygen uptake rates, *S*_*γ*_ and *S*_*ξ*_ are the drug and oxygen supply rates, and *d*_*γ*_ is the drug decay rate. In numerical implementation, we take the smaller of the cellular demand *ρ*_*γ*_/*ρ*_*ξ*_ and the current drug/oxygen level to assure that both concentrations do not fall below zero. Here, *x* represents the Cartesian coordinate system, *X*_*i*_ are the coordinates of discrete cells, *V*_*j*_ are the coordinates of discrete vessels, and *χ* is the indicator function of the local neighborhood of radius *R* around the cells *X*_*i*_ or of radius *R*_*ν*_ around the vessels *V*_*j*_, respectively:

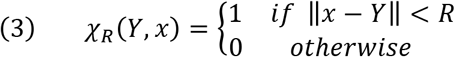

**Figure 1.**
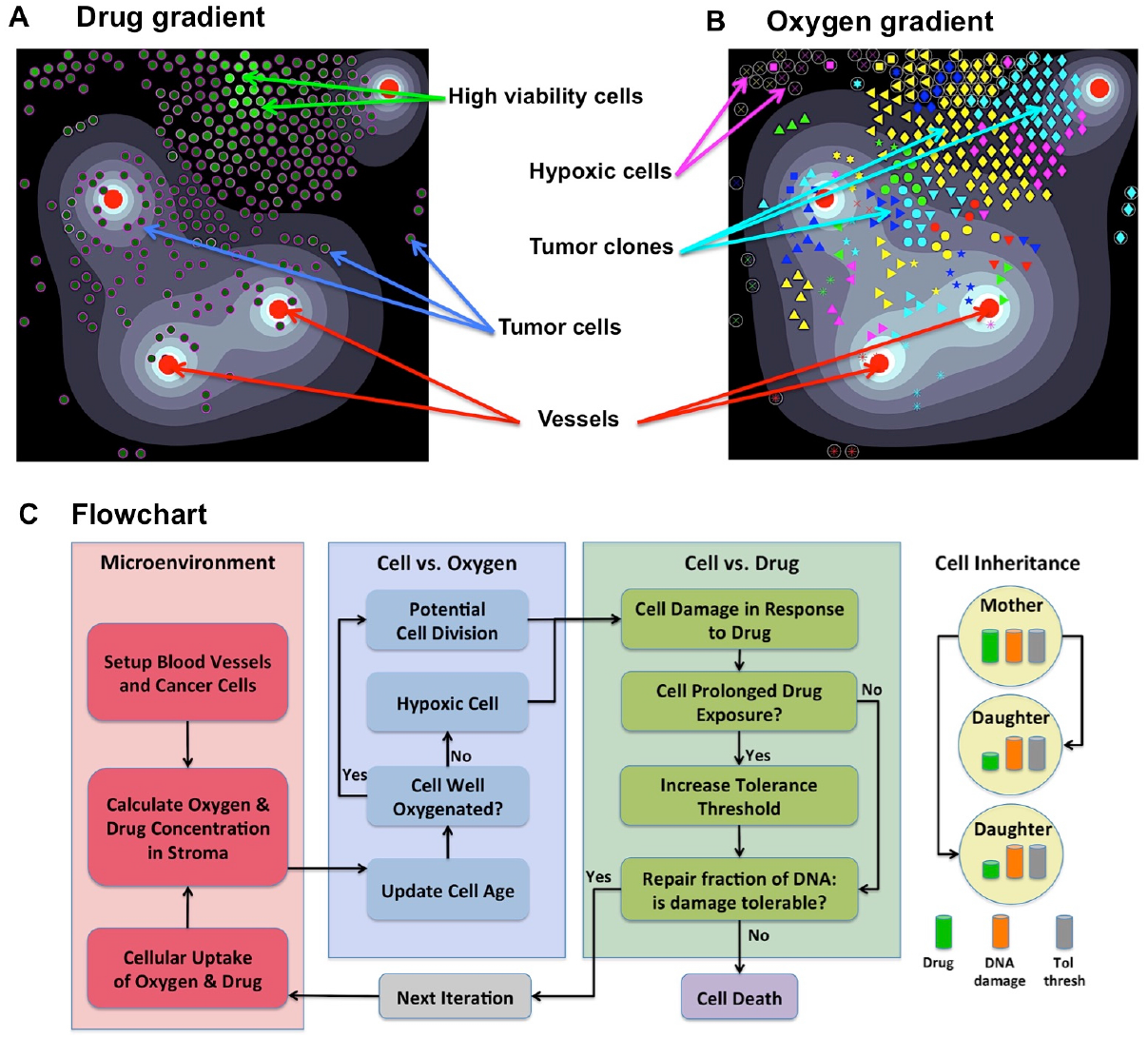
Components of the *MultiCell-LF* model. **A.** A snapshot showing an irregular drug gradient (high level-white, low level-black) and individual tumor cells color-coded according to their viability (low viability-dark green, high viability-light green). **B.** The same time snapshot showing oxygen gradient (high level-white, low level-black) and tumor clones marked by a unique symbol assigned to their initial ancestor cell (65 different symbols). Red circles in both panels represent four non-symmetrically located vessels. **C.** A flowchart showing the relationship between cell behavior and (from left to right) the tumor microenvironment; oxygen levels that regulate cell proliferation or quiescence; drug levels that regulate cell survival, adaptation or death; upon cell division daughter cells inherit from the mother cell: the damage, tolerance level and half of the accumulated drug.

The initial condition consists of a stable oxygen gradient and no drug. The sink-like boundary conditions are imposed to implicitly represent the lymphatic system.

### Individual cell dynamics

Each cell *C*_*i*_(*t*) is defined by its position *X*_*i*_(*t*), a fixed radius *R*, cell current age *A*_*i*_(*t*) and cell maturation age 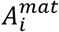. Cell division takes place upon reaching maturation age (30 hours with 5% fluctuations between the cells to avoid synchronized cell division (39, 40)) provided that the host cell is not overcrowded by other cells (14 cells within 2 cell diameters), and it is not located in the hypoxic areas (35). If the level of oxygen in a cell’s neighborhood falls below the hypoxia level (5% of vascular supply), the cell becomes quiescent and will not proliferate (flowchart in Figure 1C). Upon division of cell *C*_*i*_(*t*), two daughter cells *C*_*i*1_(*t*) and *C*_*i*2_(*t*) are created instantaneously. One daughter cell takes the coordinates of the mother cell, whereas the second daughter cell is placed near the mother cell at a random angle *θ*:

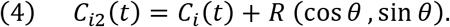

The current ages of both cells are initialized to zero, and their cell maturation ages are inherited from their mother cell with a small noise term. To preserve cell volume, the repulsive forces are introduced between overlapping cells (13, 14). Since both daughter cells are placed in a distance equal to one cell radius, the repulsive forces are exerted to push the cells apart until they reach the distance equal to cell diameter. The repulsive forces are defined as overdamped springs:

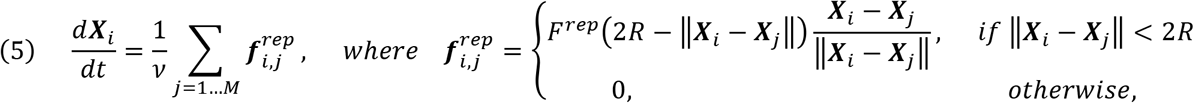

here, *ν* is the damping coefficient, *F*^*rep*^is the repulsive spring stiffness, and *2R* is the spring resting length; M denotes the number of cells that overlap with *X*_*i*_.

Upon division, both daughter cells inherit mothers’ damage level and tolerance level, while the drug absorbed by the mother cell is split into half between both daughter cells (18, 38).

### Cell resistance mechanism

Cell’s resistance mechanism is modeled as a competition between the level of drug-induced damage accumulated by the cell and the level of damage that the cell can withstand (tolerance) without committing to death. However, the cell can also adapt by increasing its tolerance level if it is exposed to the drug for a certain time (flowchart in Figure 1C, (13, 14)).

Cell damage 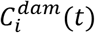 is increased proportionally to the newly absorbed amount of drug (we assume that the drug absorbed in the past has already exerted its damage effect). However, this drug-induced damage can be counterbalanced by cell natural ability for damage repair at rate *ρ*_*γ*_(three times faster that damage rate, (13, 14)):

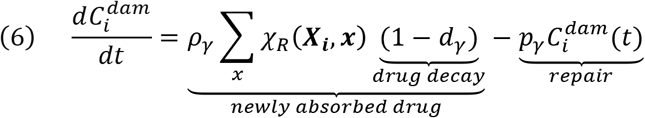

Cell exposure to high drug concentrations *γ*_*exp*_(at least 1% of the vascular supply) for a long enough time *t*_*exp*_(at least 2% of the cell cycle) results in cell adaptation to the microenvironment and in increased cell tolerance to the drug 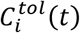 (at a slow rate of Δ_*tol*_=0.01% of the baseline tolerance value, (13, 14)):

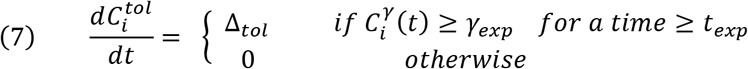

where, the amount of accumulated 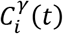 depends on its continuous absorption (at a constant rate *ρ*_*γ*_) and decay (at a rate *d*_*γ*_):

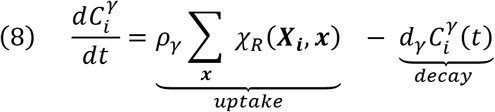

Cell death depends on whether cell damage 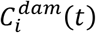 exceeds the tolerance level 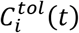. The dead cells are removed from the system. Thus, cell resistance to the drug depends on competition between the level of its damage and the level of damage the cell can withstand without committing to death.

Initially, there is a small micrometastasis consisting of 65 cells with the same baseline tolerance level, no damage, and identical proliferative properties. Each cell responds to the environmental cues, such as the level of sensed oxygen (that regulates cell quiescence or proliferation) and the amount of absorbed drug (that induces cell damage and modulates cell adaptability). The levels of drug and oxygen that the cell is exposed to during its lifetime can vary because the cell can move from one part of the tissue to another, and because drug gradient can change if the overall number of tumor cells changes. The full flowchart of cell behavior is shown in Figure 1C. During the simulation, we trace location and viability (the difference between tolerance and actual damage) of each individual cell. When the values of cell damage and tolerance steadily diverge over time, the cell is considered resistant to the drug.

### Viability trajectories of individual cells

Cell viability is defined as a difference between the level of cell tolerance to drug-induced damage and actually accumulated damage. The larger the viability value, the more non-responsive to the absorbed drug the cell is. The viability trajectory shows how the viability value is changing in time for a given cell and all its predecessors. The viability trajectory is generated backward starting from the last iteration at which the cell was alive, and going back the cells’ lifespan, the lifespan of that cell’s mother, the mother’s mother, and so on until the one of the initial 65 cells is reached (compare Figure 2B, Figure 3A-D, Figure 4B, and Figure 5).

**Figure 2.**
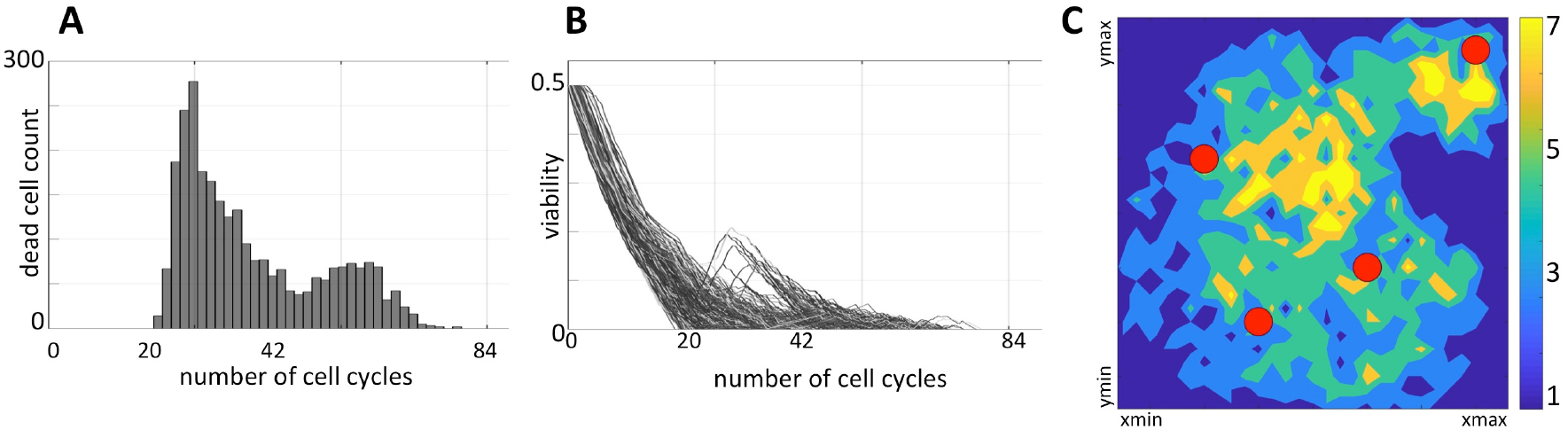
Distribution of non-resistant cells. **A.** A temporal histogram of dying cells. **B.** Individual viability trajectories for each dying cell. **C.** A density map of final locations of cells before they were annihilated by the drug-induced damage. Colors correspond to the number of cells killed at the given location during the whole simulation.

**Figure 3.**
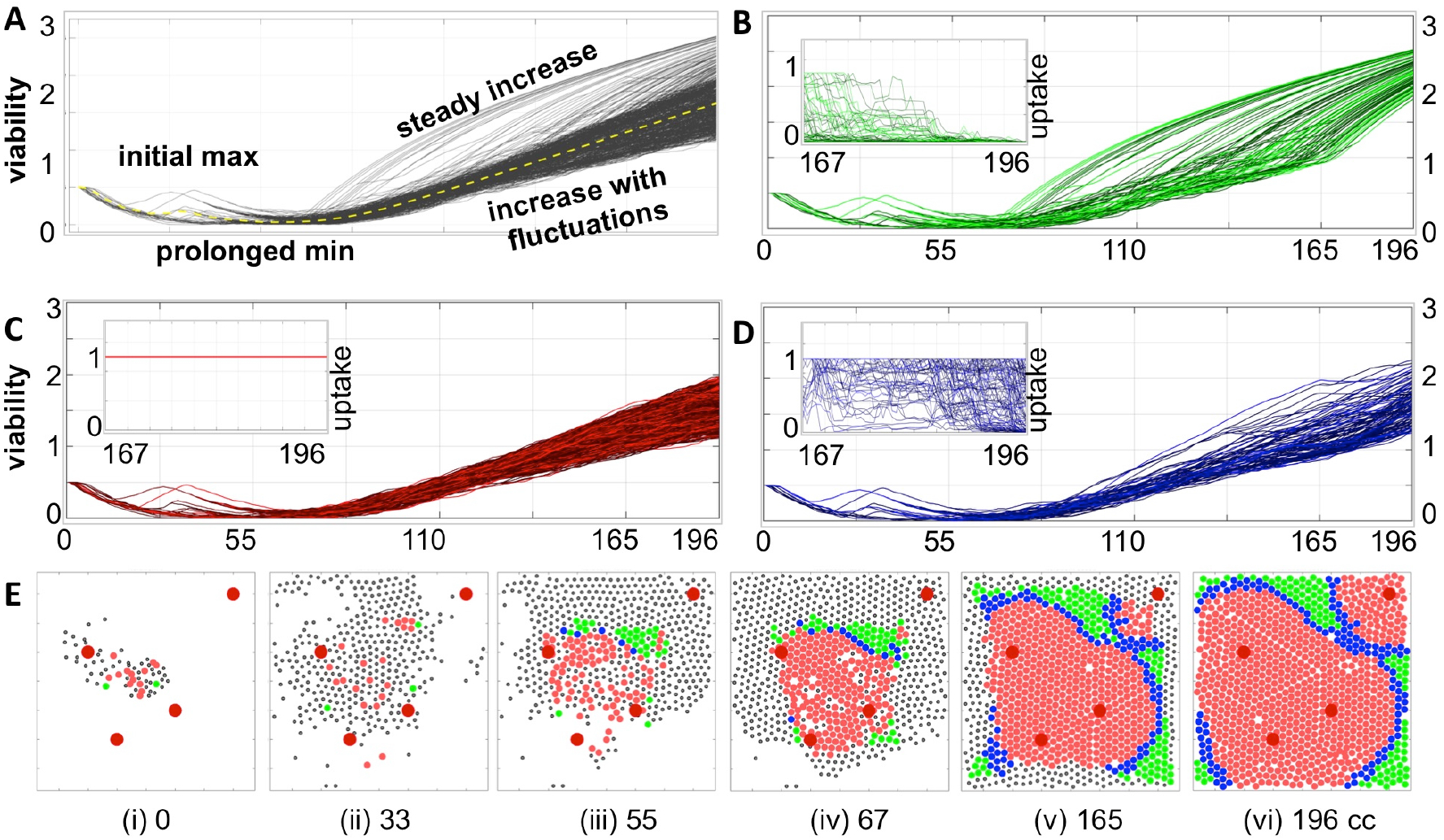
Individual cell adaptation to drug exposure. **A.** Temporal evolution of viability trajectories of all tumor cells that survived the treatment (grey lines) and the average viability value (yellow dashed line). **B**-**D.** Viability trajectories for individual cells from three distinct populations of cells with **B.** superlinear, **C.** linear, and **D.** intermediate viability patterns; the insets show drug uptake by each cell from a given subpopulation during the final period of treatment corresponding to 29 cell cycles of the simulated time: **B.** significant decrease in drug uptake, **C.** constant drug uptake, and **D.** small or short-time decrease in drug uptake. **E.** Spatial configurations of tumor cells at six different time points; colors correspond to cell subpopulations from **B**-**D**; black dots indicate cells that did not survive to the end of simulation. The larger red circles represent four vessels. All time points are shown in terms of the number of cell cycles (cc).

**Figure 4.**
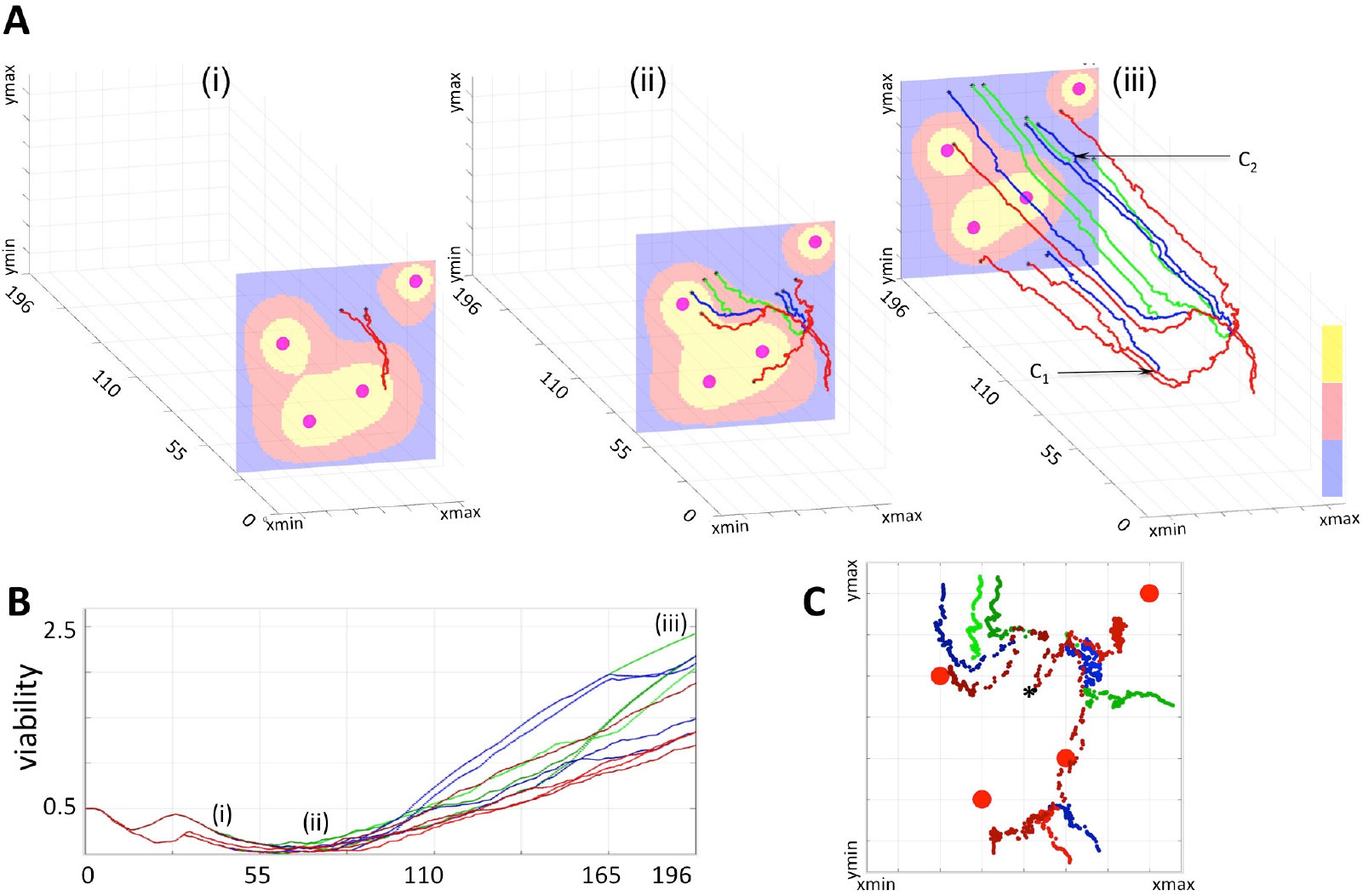
3D Spatio-temporal routes of the representative resistant cells. Twelve resistant cells with a common predecessor were traced in space and time. All line colors correspond to cell viability categories: concave (green), linear (red) and intermediate (blue). **A.** Three snapshots of routes traversed by the selected cells within a dynamically changing drug gradient in the background (high to low: yellow-red-blue; four large circles indicate the vessels). **A(i)** initial overlap in cell routes (time corresponding to 33 cell cycles, 3 cells); **A(ii)** separation of individual cell routes and significant distances traveled by the cells (about 67 cell cycles, 9 cells); **A(iii)** final map of cell routes showing very small changes in cell locations over an extended time (about 196 cell cycles, 12 cells). **B.** Viability trajectories for the selected cells. **C.** Projections on the xz plane of the individual routes for all twelve cells. A black star indicates the initial position of the common predecessor cell.

**Figure 5.**
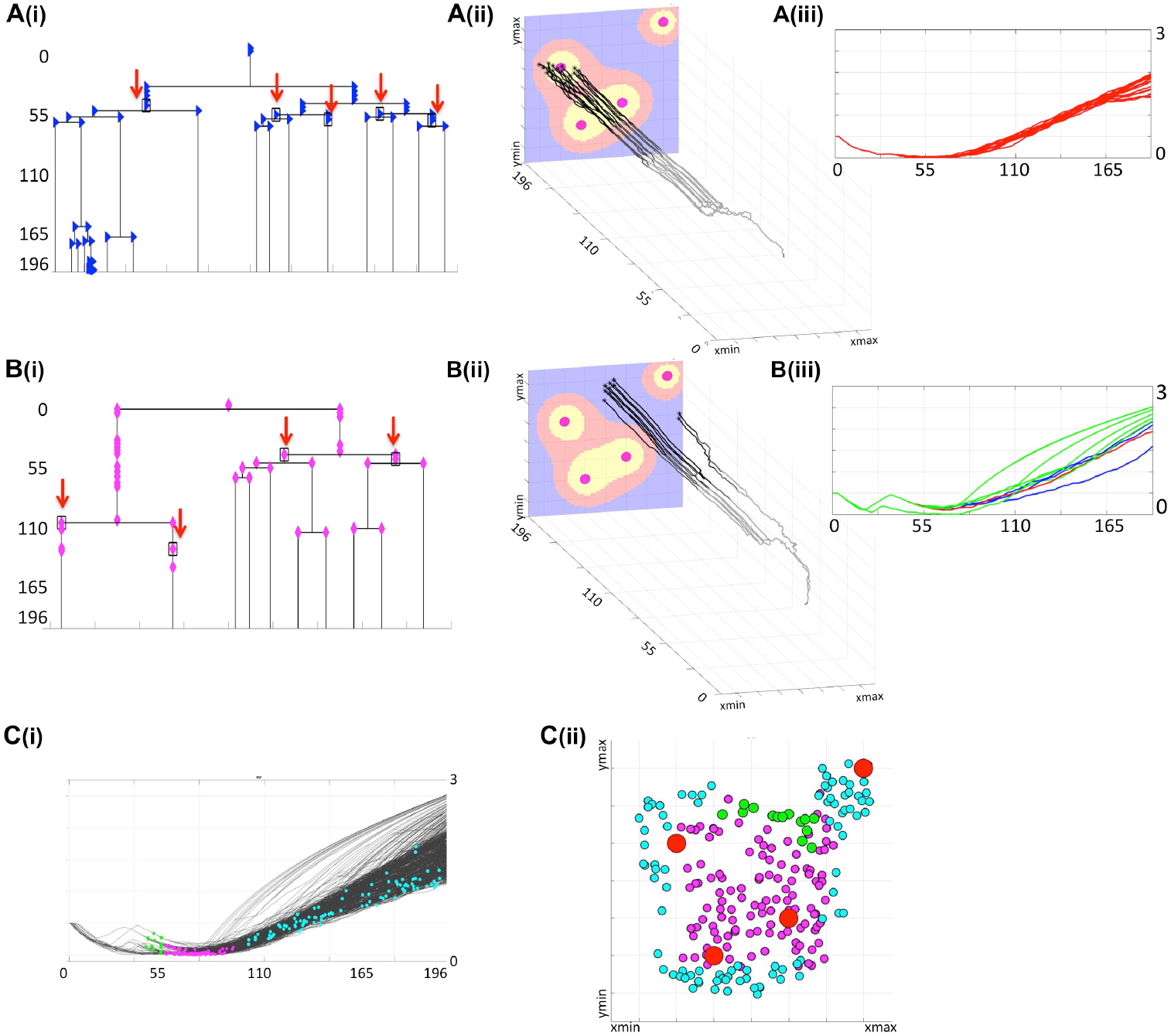
Lineage trees of survivors and cellular precursors of drug resistance. **A(i)** shows all successors of cell #6 that survived the treatment (lineage tree of survivors). The cells which have no dead successors (the precursor cells of resistance) are shown by red arrows. The corresponding spatio-temporal routes of all surviving cells that arose from cell #6 are shown in **A(ii)** and indicate changes in cells position during the treatment, and final cell locations within the tissue with a drug gradient. The corresponding viability trajectories are shown in **A(iii)**; line colors correspond to those in Figure 4. **B(i)-(iii)** shows the lineage tree of survivors, the precursor of resistance and, the 3D spatio-temporal routs, and the viability curves for all daughter cells generated by cell #26. **C(i)** shows distributions of all precursor cells along the full set of viability trajectories, with spatial localizations of all precursor cells within the tumor tissue shown in **C(ii);** colors represent time of cell birth: early (green), middle (magenta), and late (cyan); red circles show vessel locations. All time points indicate time in the number of cell cycles.

### Classification of viability trajectories and cell adaptation process

To classify how a given cell adapts to the drug exposure, we took into account both its viability trajectory and its recorded drug uptake over the last 20 cell cycles. Visually, there were two significantly distinguishable patterns: cells with constant drug uptake, and cells with viability trajectories of concave shape. Therefore, the following classification criteria were chosen: (i) the amount of absorbed drug is constant; (ii) the viability curve is monotonically increasing over at least 95% of the considered time interval (numerical second derivative of the viability curve is negative); (iii) the remaining cases. As a result, we identified: (i) a superlinear adaptation pattern (concave viability trajectories with diminished drug uptake); (ii) linear adaptation pattern (constant drug uptake and almost linear viability trajectories); (iii) intermediate pattern where cellular uptake was diminishing but the viability trajectory was not concave for the majority of time (compare Figure 3B-D and the figure insets).

### Cells 3D spatio-temporal routes/3D lineage trees

A 3D cell route shows a spatio-temporal evolutionary history of a given tumor cell; that is, it shows all recorded locations of that cells and all cell’s predecessors within the tissue patch. The 3D route is created backwards in time by linking positions (in the x-z plane, locations within a tissue at a given time) of a given cell taken at consecutive time points (y-axis) until the cell’s birth time is reached, and then repeating this procedure for all cell’s predecessors until the beginning of the simulation. The 3D spatio-temporal routes can be traced for multiple cells of the same predecessor forming a 3D spatio-temporal lineage tree (compare Figure 4A and Figure 5AB(ii)). These 3D lineage trees are an extension of classical lineage trees used to depict tumor clonal expansion. These figures synthesize information regarding cell locations, cell lineage and time in one single image.

### Lineage trees of survivors

For each of the initial 65 cells, the whole binary lineage tree can be constructed that contains all successors of this cell. A lineage tree of survivors is a subtree of the whole lineage tree and contains only these tree branches that lead to cells that survived the whole treatment (compare Figure 5AB(i) and Supplemental Figures S4-S18).

### Precursors of resistance

The precursors of resistance are these cells for which all successors survived the treatment at the end of simulation. The precursor of resistance is identified by inspecting the lineage tree generated by that cell; if the lineage tree does not contain any dead cells, its initiating cell is considered to be a precursor. We treat the cells that left the computational domain as alive, thus allow them to be successors of the precursor cells (compare Figure 5AB(i)).

## Results

We previously analyzed a parameter space of this model and identified regimes for which the whole tumor developed resistance (13, 14). Here, we summarize these results briefly. A small colony of 65 sensitive tumor cells was exposed to a drug diffusing from four irregularly placed vessels for the period of about 200 cell cycles. Initially, the tumor increased in size until the cells started responding to drug-induced damage and dying, but, finally adapting their tolerance. After about 84 cell cycles, the tumor reached a stable population. The average cell viability showed also a steady increase that confirmed the emergence of drug resistance. The evolution of tumor resistance on the population level is presented in Supplemental Figure S1. This showed that a small homogeneous cell colony exposed to a drug gradient can acquire resistance. The final tumor contained the offsprings of 15 initial cells only; the successors of the remaining 50 initial cells went extinct. The rest of the paper is devoted to analysis of resistance at the individual cell level, whether spatial structure of the microenvironment play a role in the emergence of resistance, and which cell lineages drive resistance of the whole tumor.

### Temporal distributions of dying cells confirms the drug-induced resistance

The fate of each cell depends on both the accumulated damage and the level of damage that the cell can withstand without committing to death. To determine how the resistance is acquired in individual cells, we need to understand the conditions leading to cell death. Initially, each cell has some baseline tolerance level and no accumulated damage. With time, the absorbed drug induces damage to the cell, while the cell can also adapt to the surrounding extracellular conditions that leads to increase in its tolerance. The cell dies when the level of cell damage exceeds the level of cell tolerance to damage. During the simulated treatment, the initially sensitive cells either develop resistance or respond to the treatment and die. In fact, about 75% of the initial 65 cells did not produce offsprings that were able to survive to the end of the treatment period. Since some cells located near the domain boundaries might have been pushed outside of the observed tissue region by the pressure from their growing neighbors, these cells are assumed to move to the other tissue areas and are removed from our system. Here, we only consider cells that remained inside the tissue domain until they were annihilated by the drug. The summary of spatial and temporal analysis of dying cells is shown in Figure 2.

During the initial period of treatment, no cells were dying (no cell counts in histogram in Figure 2A) since they must accumulate the drug-induced damage to overcome the baseline tolerance level. However, the viability trajectories for numerous cells decreased during this time (Figure 2B) confirming that these cells were accumulating damage. Each curve in this graph corresponds to one cell and traces in time the viability values of this cell and all its predecessors, back to one of the initial 65 cells. This period corresponds to a steady tumor growth shown in Supplemental Figure S1(i-ii). The first peak in cell death histogram and a time interval when cell viability trajectories reached zero match the significant reduction in the overall tumor size (Supplemental Figure S1(iii)). The second peak in the death histogram is much smaller since a large number of cells have already developed resistance and only a small subpopulation of cells remained still sensitive to the drug (compare to steady tumor growth in Supplemental Figure S1(iv-v)). After the time corresponding to about 84 simulated cell cycles, no more cells have died. Similarly, all viability trajectories for these dying cells reached the zero value at or before this time (Figure 2B). This confirms that all remaining tumor cells in the observable tissue patch have developed a drug-induced resistance. Spatially, the tissue regions that are most prone to cell death are situated either near the single vessel in the top-right corner or in the region near the tissue center between the remaining three vessels (Figure 2C). It is worth noting, that in our previous work (13), we identified the model parameter regimes for which the tumors got extinct, thus the development of drug-induced resistance is not an intrinsic property of our model.

### Cell adaptation to drug exposure can progress in three distinct ways

To determine how individual cells contributed to the overall tumor resistance, we analyzed the viability trajectories of each cell that survived the treatment (Figure 3A). These graphs confirm our previous observations of several phases in the evolution of resistance in the individual tumor cells: from initial identical viability values, to viability decrease due to the damage being accumulated, to a transient increase in viability when the mechanism of tolerance became activated (initial max), to prolonged reduction in viability values due to accumulated damage approaching the individual cell tolerance level leading to cells adaptation to the drug (prolonged min), to a continuous increase in cell viability when the tolerance mechanism gains a lead. Despite the fact that all surviving cells originated from identical predecessor cells and that they shared very similar viability trajectories for the first 55 cell cycles, we identified three patterns of cell adaptation that resulted in drug-induced resistance (Figure 3B-D).

The first cell subpopulation is characterized by rapid increase in viability values that form concave curves of distinct durations (Figure 3B). In all these cases, there is also a reduced absorption of the drug for a significant length of time (at least 29 cell cycles, inset in Figure 3B). The diminished drug uptake is a result of drug concentration being below the cell’s demand. A closer analysis of cell spatial distributions over time shows that this subpopulation occupied tissue regions distanced from the vessels and, more importantly, was surrounded by other cells (green circles in Figure 3E(i-vi)). This was a combined effect of cells’ proliferation and their passive relocation due to physical pressure from other growing cells. Since the cells remained in the areas poorly penetrated by the drug for a prolonged time, it resulted in rapidly increasing cell viability that is manifested by the concave (“rainbow-like”) shape of the viability curves. The second subpopulation consists of cells with nearly linear increase in viability values (Figure 3C) and with constantly high drug absorption (inset in Figure 3C). The early predecessors were located in between the three central vessels (red circles in Figure 3E(ii-iii)), and thus were exposed to moderate drug concentrations. This resulted in faster increase of drug-induced tolerance than drug-induced damage and in the steady increase in cell viability values. The fluctuations in almost linear viability patterns arose from competition between gained tolerance, acquired damage and damage repair. This led to repopulation of the space between the blood vessels (Figure 3E(iv)). Subsequently, the cells were able to survive in the areas well penetrated by the drug, even in the vicinity of the blood vessels (Figure 3E(iv-vi)). The remaining resistant cells manifest an intermediate behavior with regards to drug absorption, as it decreases over a very short time near the end of the treatment period (Figure 3D) but not as pronounced as in the subpopulations with concave viability curves. This subpopulation also acquired a quite distinct spatial pattern on a border between two other subpopulations (Figure 3E(iii-vi)). These cells are transient in the sense that their characteristics may change during the treatment. For example, a cell with linear viability values may become transient if it gets surrounded by other cells, and becomes protected from drug exposure (cell C_1_ in Figure 4A(iii)). Similarly, a transient cell can give birth to a cell that falls into the category of concave viability if it moves to the poorly penetrated area (cell C_2_ in Figure 4A(iii)).

### The 3D cellular routes delineate spatio-temporal dynamics of cell adaptation

To more closely examine how cells from all three categories can adapt to the treatment, we selected twelve cells (four from each category) with one common predecessor (Figure 4) and traced their locations within the tissue during the whole simulation.

The 3D spatio-temporal routes traversed by each cell are shown in Figure 4A at three different time points together with the drug profile at that time. Here, the xz-plane represents cell positions within the tissue, and y-axis represents the time. Note, that the drug distribution profile at each time point is different despite the continuous drug influx from the vessels because drug absorption depends on the total number of cells in the tissue, and this cell number varies in time. The presented exemplary cells were chosen intentionally to show a variety of spatial and temporal dynamics that may lead to cell survival, adaptation and acquired resistance. The corresponding twelve viability trajectories are presented in Figure 4B to confirm characteristics of each cell. Since these cells have a common predecessor, there is a period of time when both the viability trajectories and the 3D routes overlap and thus the number of observable curves is limited (Figure 4AB(i)). However, these curves eventually split up in both figures into eight separate lines (Figure 4AB(ii)). Furthermore, individual cells were able to move at significant distances from the position of their common predecessor (Figure 4A(ii) and Figure 4C). This was due to the pressure imposed by other growing cells. From this point on, the viability trajectories steadily increased (Figure 4B(iii)), but cells’ routes deviated only insignificantly forming almost horizontal lines (Figure 4A(iii)). This was due to cell overcrowding by numerous neighbors that resulted in cells’ prolonged dormancy, without division. This ultimately contributed towards cell survival and steady increase in cell viability. We intentionally selected a case in which the initial predecessor cell was able to give rise to successors from each of the three categories. However, out of 15 initial cells which successors survived the whole treatment, four generated cells in all three categories, three produced cells in two categories and eight gave rise to cells in a single category.

### Lineage tree analysis identifies the cells that drive resistance

Less than a quarter of cells that formed the initial micrometastasis (15 out of 65) produced successors that survived the whole chemotherapeutic treatment. Here we examined the lineages of each subpopulation in order to identify how drug resistance developed for each of them. We inspected the full lineage trees for each of the survived subpopulation and identified subtrees containing only those branches that led from the initial cell to cells that survived the whole treatment. The branches leading to dead cells were omitted. If one of the daughter cells left the domain, but the other survived, its symbol was indicated along the vertical line connecting that cell with its mother cell. These structures represent the lineage trees of survivors. Two representative examples generated by the initial cells with indices #6 and #26 are shown in Figure 5AB(i). The corresponding spatio-temporal routes traversed by these cells are shown in Figure 5AB(ii). Additional snapshots of spatio-temporal routes at different time points are shown in Supplemental Figure S2 and Figure S3. The cell viability trajectories are shown in Figure 5AB(iii). These examples illustrate different cases of cellular adaptation observable among all survived subpopulations. The cells for which viability increases linearly are located in well-penetrated areas. These cells were able to survive the drug insult for a prolonged time since they were surrounded by other cells that absorbed the drug creating a protective niche (Figure 5A(ii-iii)). Cells with concave viability trajectories are located in poorly penetrated areas, often equidistant from the vessels, where damage induced by the drug is lower that the ability of the cell to repair damage (Figure 5B(ii-iii)). For some lineage trees of survivors, their spatio-temporal routes may have multiple spatially separated branches due to the proliferation and pressure from neighboring cells. In other cases, the routes do not deviate significantly in space and form horizontal lines. This is due to overcrowding that limits cell proliferation and migration (other 3D routs are discussed in Suppl Figures S4-S18).

For each lineage tree of survivors, we identified the subtrees that do not contain any dead cells; that is, all branches of these subtrees point either to cells that survived the whole treatment or to cells that left the domain (these cells have positive viability values, so they are alive). The roots of such subtrees are considered to be the precursors of drug resistance, since none of their successors underwent drug-induced death. The precursor cells are indicated by black rectangles and red arrows in the trees shown in Figure 5AB(i) (for clarity, the branches leading to the cells that left the domain are omitted from the graphs). In total, there were 224 precursor cells emerging from all 15 lineage trees of survivors. They all are pictured in Figure 5C(i) along the viability trajectories to indicate the time at which they emerge. In Figure 5C(ii), these cells are cumulatively projected on the tissue space to show the initial locations of the precursor cells.

The cell colors correspond to a time period at which they first appeared. The very first precursor cells have arisen in the area poorly penetrated by the drug between the single vessel in the top-right corner, and the three other vessels (cells shown in green in Figure 5C). Such areas are known as drug sanctuaries or refugia. The next cohort of precursor cells emerged in the center of the tissue between the three blood vessels (indicated by magenta dots in Figure 5C). While, in principle, these areas can be better penetrated by the drug, they actually form protective niches (protectorates) in which the precursor cells may be shielded from the exposure to the drug by the surrounding cells. The final cohort of precursor cells (indicated by cyan dots in Figure 5C) was emerging over a longer period of time and mostly in the areas located closer to the tissue boundaries in the hypoxic or nearly-hypoxic niches. Interestingly, none of the precursor cells were located directly at the concave viability trajectories. This indicates that all precursor cells emerged as a result of a direct competition between drug-induced cell damage and acquired tolerance, and that the increase in cell viability was amplified (in fast superlinear fashion) in cells that have already developed resistance.

## Discussion

We presented here a study analyzing how resistant cell lineages arise in micrometastases exposed to a systemic chemotherapeutic treatment. This research is an extension of our previous work (13, 14) that focused on the emergence of drug-induced resistance on a cell population level. While we followed the previous mathematical model setup and considered a small tumor growing in a heterogeneous microenvironment, the individual-cell perspective and novel evaluation methods allowed us to identify a new microenvironmental niche prone to the emergence of resistant cells. In addition to previously reported *refugia* characterized by low drug penetration due to their distance from the vasculature and the *hypoxic or near-hypoxic niches* in which cells were able to thrive and repair the drug-induced damage, we also located areas in which cells were not exposed to lethal drug concentrations because they were shielded by other cells absorbing the excess of the drug—the *protectorates*. We also recognized that certain cells gave rise to lineages of resistant cells (precursors of resistance) and correlated three temporal periods with three different spatial locations at which such cells emerged. This supports the hypothesis that tumor micrometastases do not need to harbor cell populations with pre-existing resistance, but that individual tumor cells can adapt and develop resistance induced by the drug during the treatment.

The novel analysis and visualization methods developed here, such as the lineage trees of survivors, the method to identify the precursors of resistance and the 3D sptatio-temporal evolution routes, can enhance the library of tools used with other hybrid mathematical models (41–43).

Moreover, we showed that once the cells have developed resistance, they were able to elevate their viability either in a fast superlinear manner or in a slower, linear fashion, depending whether they moved toward the refugia areas or not; a small population of transient cells that could transfer from the linear to superlinear populations was also observable. This is in line with the theory of mixed models of tumor evolution (28), in which different evolution forms can occur in parallel or can shift from one form to another as a result of changes in tumor size or due to microenvironmental selection forces.

Our results can be also placed within a context of tumor ecology (44, 45), such as the ecological concepts of microenvironmental niche partitioning and niche construction (46). In the former case, different cell subpopulations are driven into distinct tissue compartments by the microenvironmental selection forces—we observed that certain cell subpopulations were harbored within the refugia areas or within the hypoxic niches. In the latter, the cells are able to modify their own surroundings to create a favorable microenvironment—we observed the formation of cellular protectorates characterized by microenvironmental conditions distinct from the surrounding areas. This spatial heterogeneity in tumor microenvironments is often referred as ecological habitats (47, 48) that can lead to unique fitness landscapes and selection for different cell phenotypes and genotypes, even under the same extrinsic pressure such as anti-cancer therapy. Our simulations showed that individual cell viability was changing over time that encourages revisiting the idea of a static fitness landscape, and supports the view that cell fitness is not a constant value, but a function of the environmental context (46, 49). While we did not explicitly model any genetic mutations, the observable changes in tumor cell viability could be potentially linked to changes in cell gene expression.

Ultimately, the link between ecological changes within the tumor microenvironment and tumor evolutionary changes will reflect on patients’ clinical outcome. While the systemic chemotherapy is often used in the clinical protocol in order to minimize the tumor metastatic spread, it should be taken into account that such therapy may stimulate progression of the nearly-killed cells toward resistance. Therefore, the approaches targeting the resistance-inducing strategies may prove more effective than targeting the tumor cells directly. This is similar in concept to eco-evo drugs from the field of microbial antibiotic resistance (50) Some such preconditioning mechanisms have been tested in cancer cells and already showed promise (20, 51, 52); however, more research in this area is needed.

## Acknowledgment

This work was supported in part by the U01-CA202229 Physical Sciences Oncology Project (PS-OP) grant from the US National Institutes of Health (to KAR). JPV thanks the "Global Challenges for Women in Math Sciences” program of the Mathematics Faculty of the Technical University of Munich for the Entrepreneurial Award to further this collaborative project.

