## Supplemental Material for "Drug-induced resistance in micrometastases: analysis of spatio-temporal cell lineages"

#### Analysis of tumor resistance on the cell population level

The development of drug-induced resistance proceeded in five stages detailed in Figure S1.

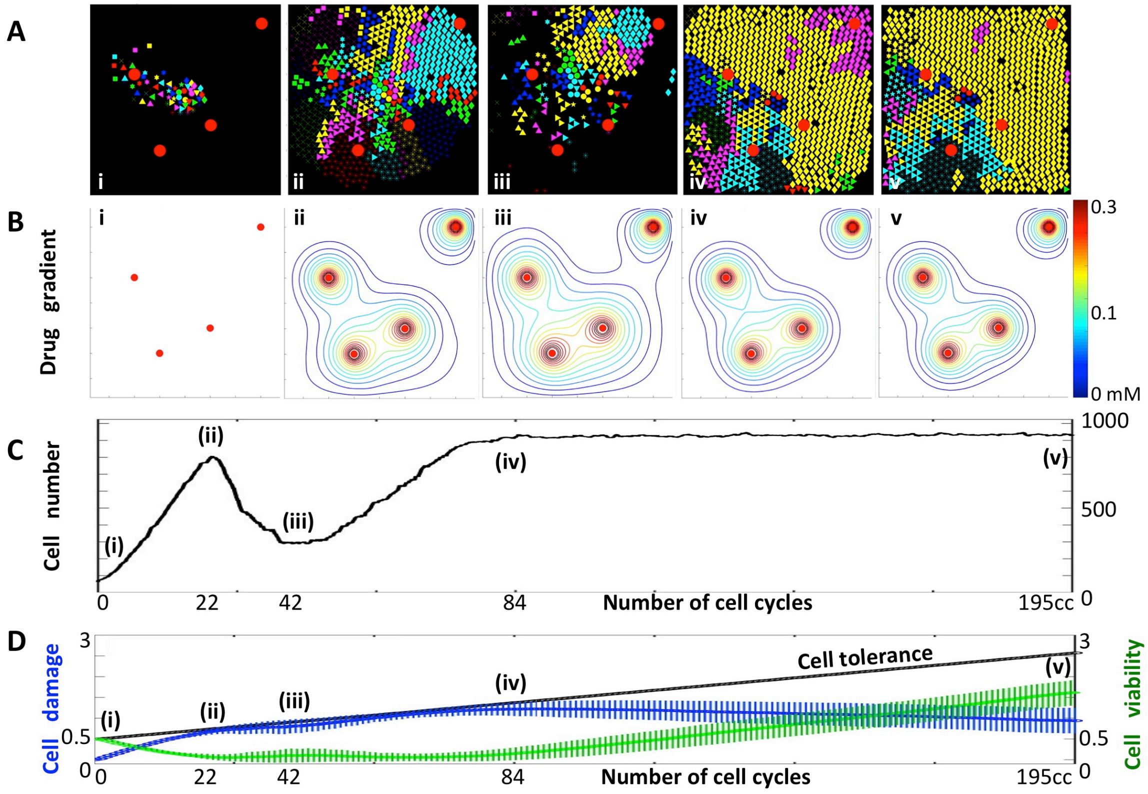


**Figure S1. Development of a tumor with drug-induced resistance. A.** Representative snapshots showing spatial configuration of tumor cells colored according to their clone of origin (65 initial cells) at 5 specific time points of tumor development. **B.** The corresponding spatial distributions of the drug. Numbers (i)-(v) indicate specific time points in tumor development: (i) initiation; (ii) peak in the initial tumor growth; (iii) tumor regression due to drug cytotoxicity; (iv) tumor expansion due to acquired resistance; (v) the stable resistant tumor. **C.** The change in the total number of tumor cells over the period of 195 cell cycles from 65 to ~900 cells. **D.** The evolution of average population-level metrics with standard deviation values represented as vertical lines. Average cell viability from initial 0.5 to maximal value around 2 (arbitrary units, a.u) is shown as a green line; average cell damage which starts at 0 (a.u) and reaches 2 (a.u) at the peak is shown in blue; and average cell tolerance level initiated at 0.5 (a.u) and reaching 3 (a.u) is shown in black.

A small micrometastatic colony consisting of 65 tumor cells was exposed to a drug diffusing from four irregularly placed vessels (Figure S1A(i)). The temporarily and spatially changing drug concentration influenced tumor development over the period of about 200 cell cycles resulting in drug-induced resistance. Initially, all cells had the same sensitivity to the drug and differed only in their locations within the tissue. With time, these cells became exposed to a changing concentration of the drug (Figure S1B), and they were responding by acquiring drug-induced resistance and by adapting their tolerance (Figure S1D). Since cells at different locations experienced drug changes differently, the viability of individual cells was evolving with different dynamics (which is indicated by the increasing standard deviations of average viability values in Figure S1D). After the time corresponding to about 84 cell cycles, the tumor stopped responding to the drug and reached a stable population of about 900 cells (Figure S1C) that filled the whole observed tissue patch (Figure S1A). The average cell viability showed also a steady increase that confirmed the emergence of drug resistance (Figure S1C). This indicated that the tumor as a whole became resistant to the drug. The final tumor contained the offsprings of 15 initial cells only; the successors of the remaining 50 initial cells went extinct (Figure S1A).

#### Evolution of spatio-temporal routes for selected initial cells

A 3D spatio-temporal evolutionary history graph synthesizes information about cell lineage with cell location changes over time, and the temporal and spatial changes in tumor microenvironment.

##### Supplementary Figure S2

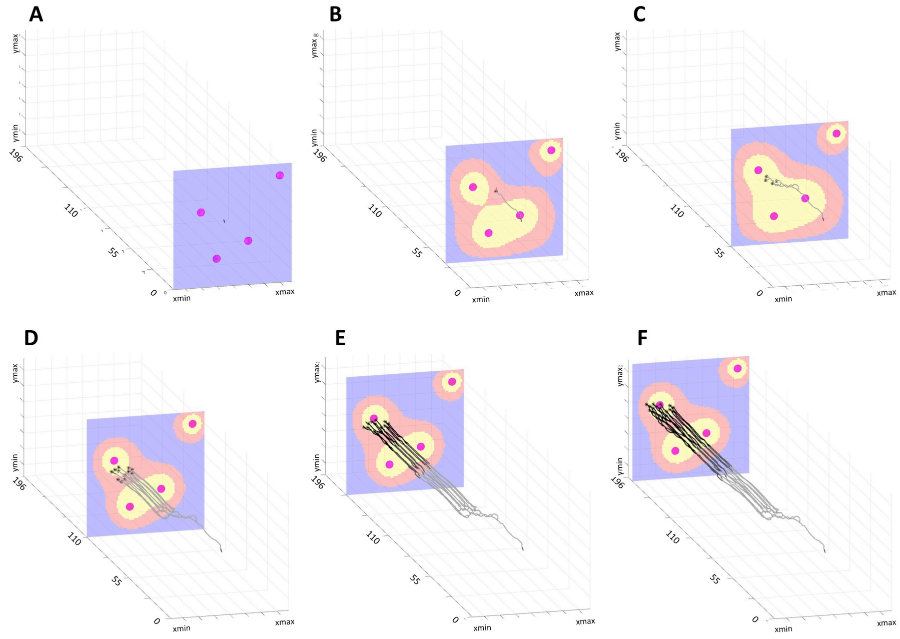


**Figure S2.** A sequence of 3D spatio-temporal routes of all surviving cells initiated from cell #6 at times corresponding to: **A.** treatment initiation, **B.** 30 cell cycles (cc), **C.** 55 cc, **D.** 110 cc, **E.** 165 cc, and **F.** 196 cell cycles. Background images show the drug gradient at a given time (levels high-to-low: yellow-red-blue).

Since the drug supply from four irregularly spaced vessels (red circles) is continuous, the changes in tumor microenvironment depend on the number of cells that absorb the drug. Initial changes in cell position within the tissue depend on its relocation to the areas that are deprived of the drug and allow the cell to develop resistance. The almost horizontal cell routes are the result of cell overcrowding.

##### Supplementary Figure S3

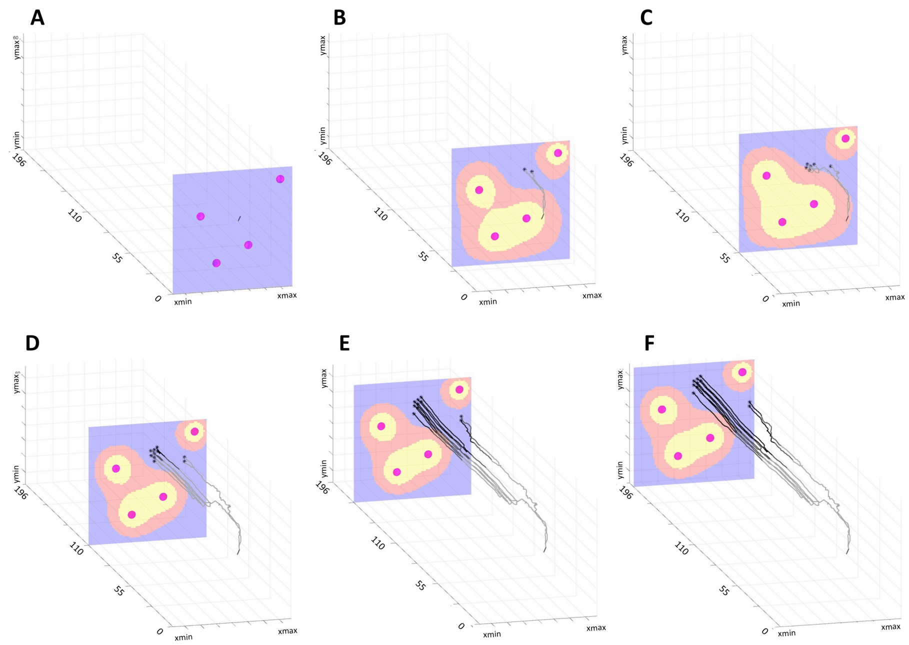


**Figure S3.** A sequence of 3D spatio-temporal routes of all surviving cells initiated from cell #26 at times corresponding to: **A.** treatment initiation, **B.** 30 cell cycles (cc), **C.** 55 cc, **D.** 110 cc, **E.** 165 cc, and **F.** 196 cell cycles. Background images show the drug gradient at a given time (levels high-to-low: yellow-red-blue).

#### The lineage tree analysis for individual initial cells

For the initial cells cluster, only a fraction of cells was able to produce offsprings that survived until the end of the computational experiment. In the discussed example, only 15 out of 65 initial cells produced survivors and generated the linage tree of survivors. For the remaining initial cells, all their offsprings died from drug-induced damage. For each of the 15 successful cells, the full linage tree, the linage tree of survivors and the 3D spatio-temporal lineage tree are shown in Supplemental Figures S2-S16.

The *full lineage tree* is a binary tree with the initial cell at its root (showed as the top cell) and all daughter cells linked to their mother cells by vertical lines. The cells that survived (leaves of the tree) have vertical lines extended to the bottom of the graph. The whole cell sequences that lead from the initial cell to the tree leaves are circled. These cells form the *lineage tree of survivors*. In the lineage tree of survivors, the cells on the same vertical line represent cells for which one of the daughters died or was pushed out of the simulations domain, and only one daughter cell survived (thus the three does not branch). For all cells from the tree of survivors, their positions within the tissue domain taken over the whole simulated time are pictured in the *3D lineage tree*. Only the final spatial distribution of the drug is shown in this graph.

#### Supplementary Figure S4

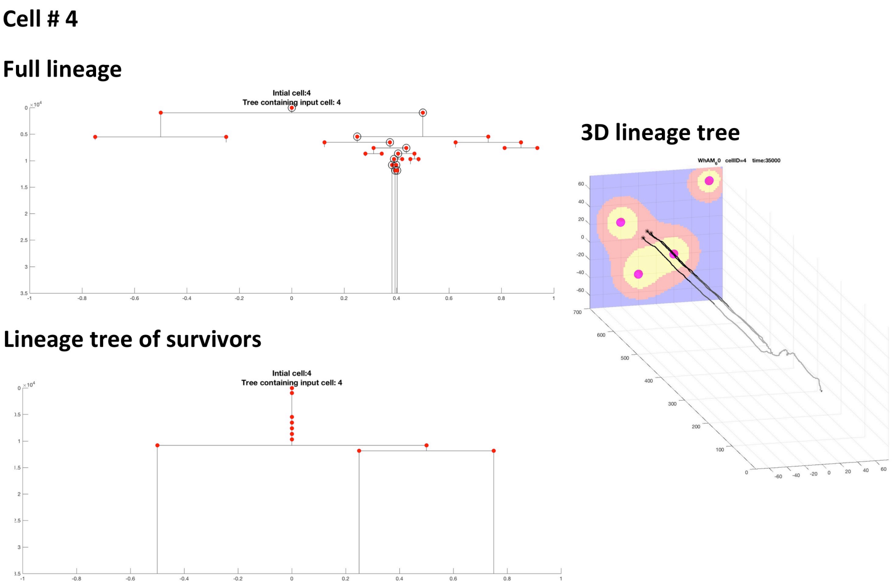


**Figure S4.** The full lineage tree for the initial cell #4 (top left), the lineage tree of survivors (bottom left), and the 3D spatio-temporal routes of all surviving cells initiated from cell #4.

#### Supplementary Figure S5

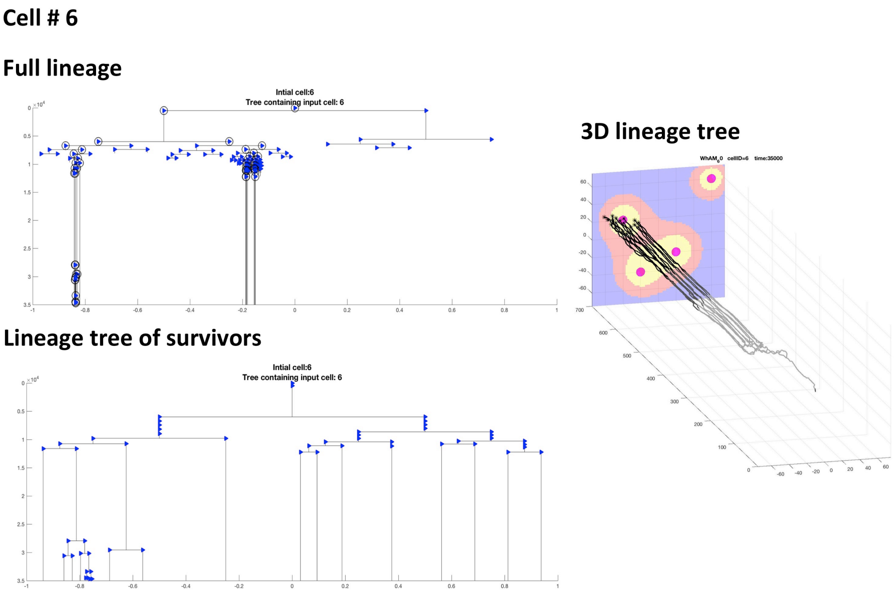


**Figure S5.** The full lineage tree for the initial cell #6 (top left), the lineage tree of survivors (bottom left), and the 3D spatio-temporal routes of all surviving cells initiated from cell #6.

#### Supplementary Figure S6

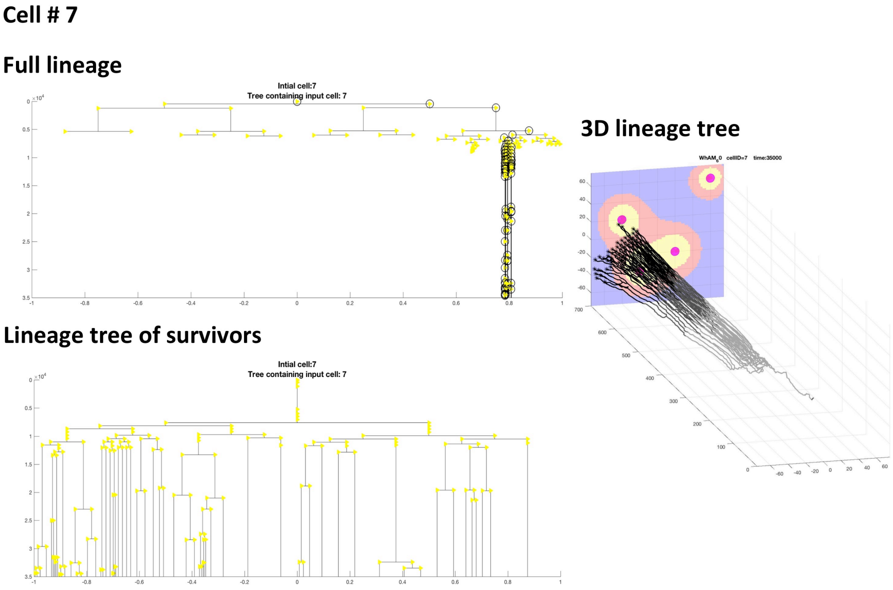


**Figure S6.** The full lineage tree for the initial cell #7 (top left), the lineage tree of survivors (bottom left), and the 3D spatio-temporal routes of all surviving cells initiated from cell #7.

#### Supplementary Figure S7

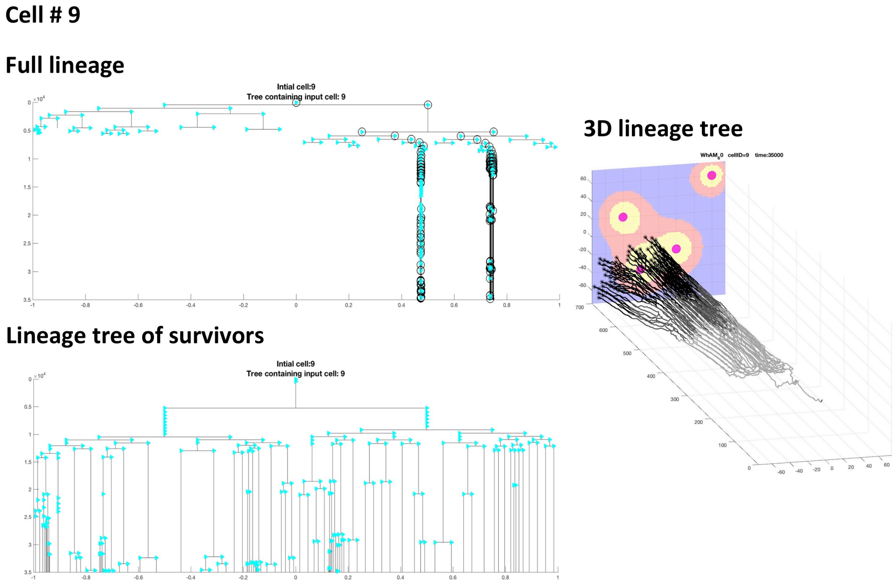


**Figure S7.** The full lineage tree for the initial cell #9 (top left), the lineage tree of survivors (bottom left), and the 3D spatio-temporal routes of all surviving cells initiated from cell #9.

### Supplementary Figure S8

##### Cell # 11

##### Full lineage

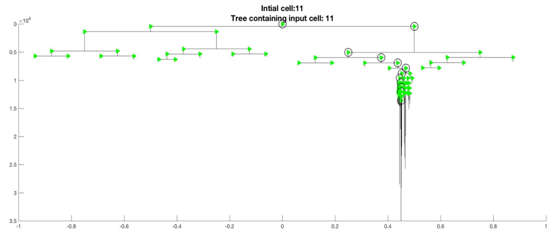


##### 3D lineage tree

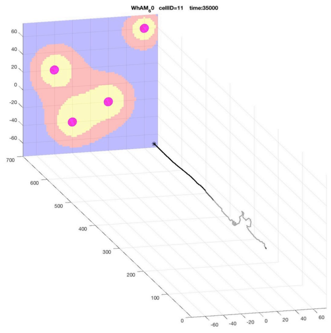

##### Lineage tree of survivors

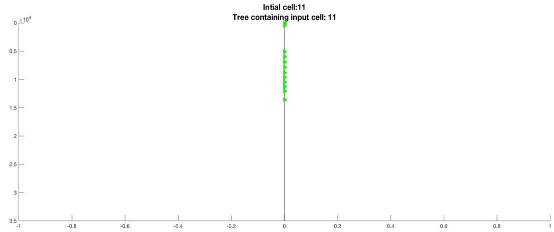


**Figure S8.** The full lineage tree for the initial cell #11 (top left), the lineage tree of survivors (bottom left), and the 3D spatio-temporal routes of all survived cells initiated from cell #11.

### Supplementary Figure S9

##### Cell # 18

##### Full lineage

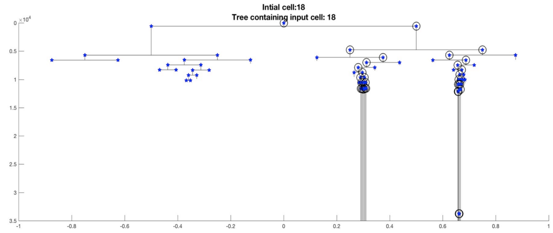


##### 3D lineage tree

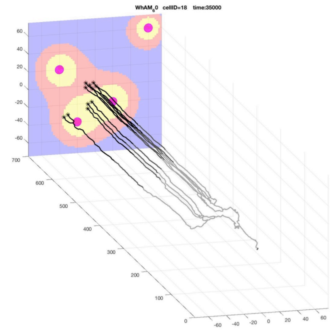

##### Lineage tree of survivors

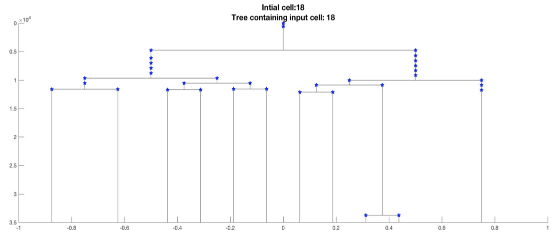


**Figure S9.** The full lineage tree for the initial cell #18 (top left), the lineage tree of survivors (bottom left), and the 3D spatio-temporal routes of all survived cells initiated from cell #18.

#### Supplementary Figure S10

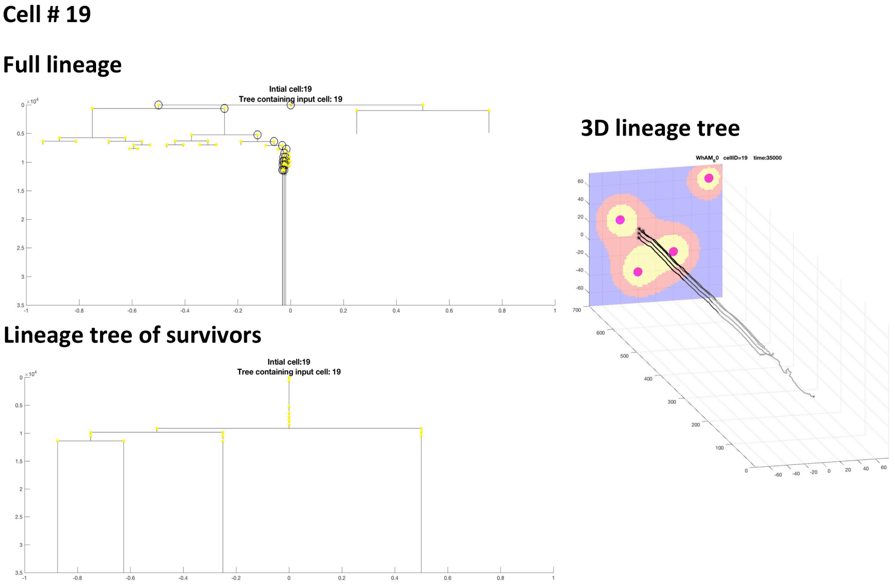


**Figure S10.** The full lineage tree for the initial cell #19 (top left), the lineage tree of survivors (bottom left), and the 3D spatio-temporal routes of all survived cells initiated from cell #19.

#### Supplementary Figure S11

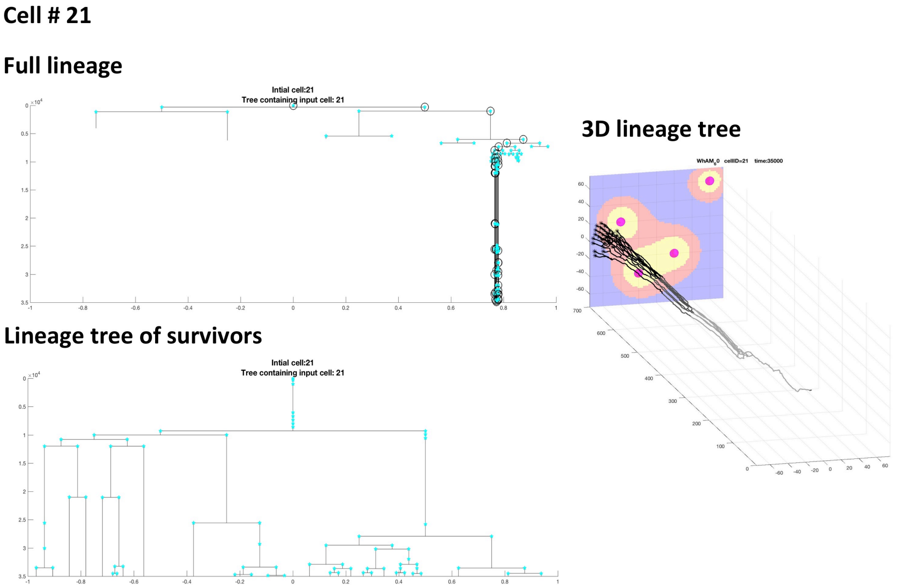


**Figure S11.** The full lineage tree for the initial cell #21 (top left), the lineage tree of survivors (bottom left), and the 3D spatio-temporal routes of all survived cells initiated from cell #21.

#### Supplementary Figure S12

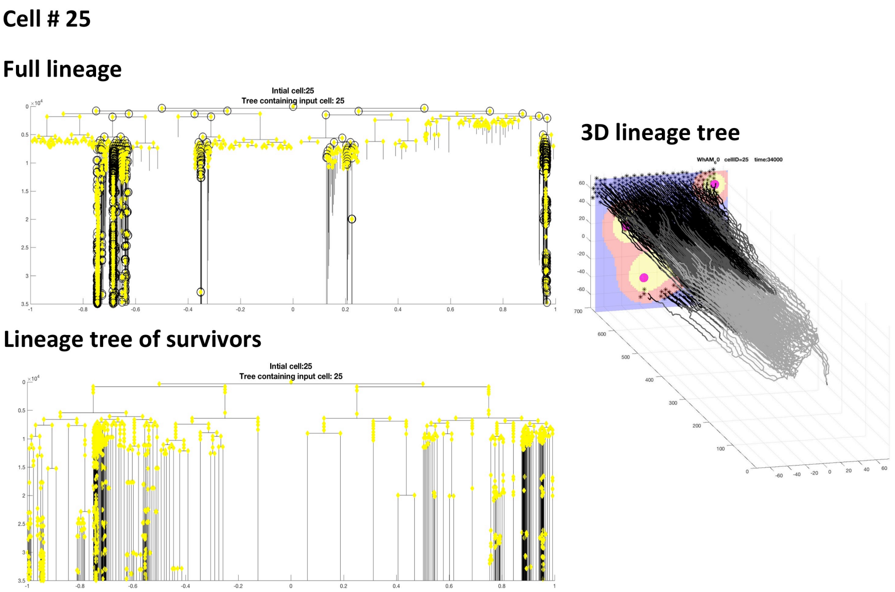


**Figure S12.** The full lineage tree for the initial cell #25 (top left), the lineage tree of survivors (bottom left), and the 3D spatio-temporal routes of all survived cells initiated from cell #25.

#### Supplementary Figure S13

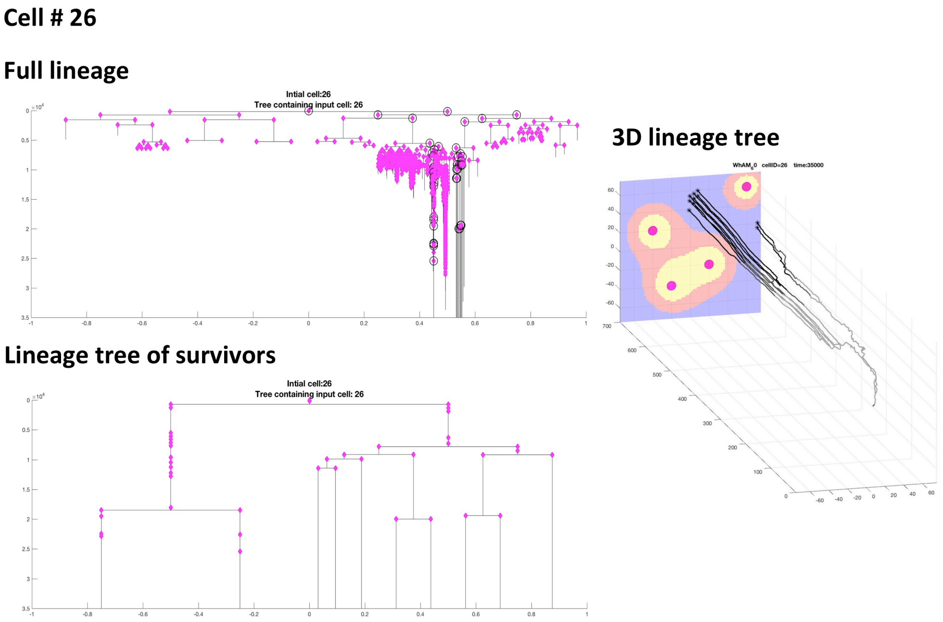


**Figure S13.** The full lineage tree for the initial cell #26 (top left), the lineage tree of survivors (bottom left), and the 3D spatio-temporal routes of all survived cells initiated from cell #26.

### Supplementary Figure S14

**Cell # 33**

**Full lineage**

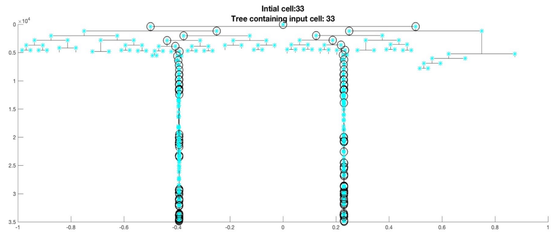


**Lineage tree of survivors**

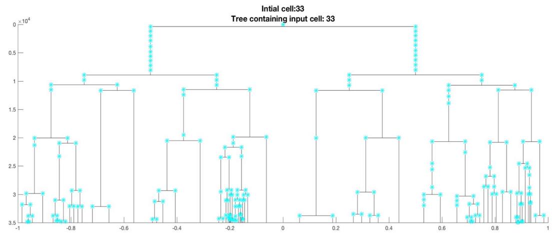


**3D lineage tree**

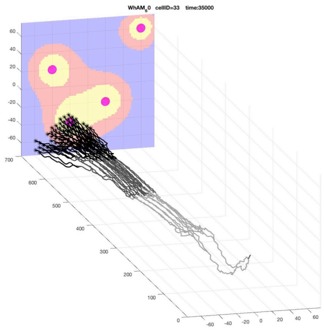


**Figure S14.** The full lineage tree for the initial cell #33 (top left), the lineage tree of survivors (bottom left), and the 3D spatio-temporal routes of all survived cells initiated from cell #33.

### Supplementary Figure S15

**Cell # 38**

**Full lineage**

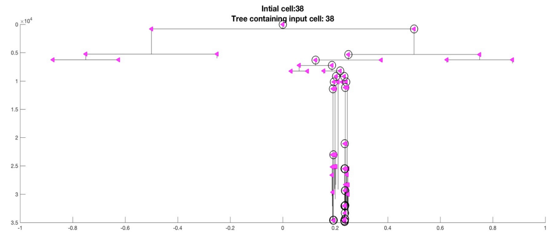


**Lineage tree of survivors**

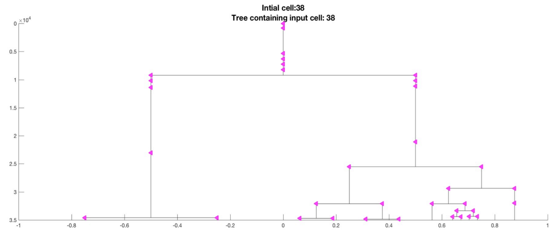


**3D lineage tree**

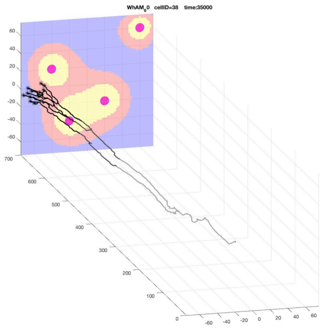


**Figure S15.** The full lineage tree for the initial cell #38 (top left), the lineage tree of survivors (bottom left), and the 3D spatio-temporal routes of all survived cells initiated from cell #38.

### Supplementary Figure S16

##### Cell # 41

##### Full lineage

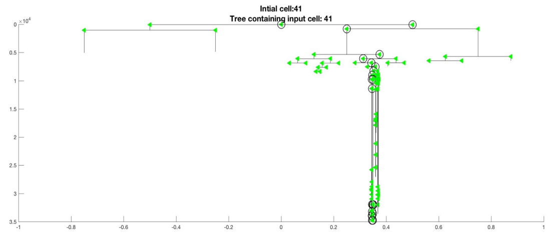


##### 3D lineage tree

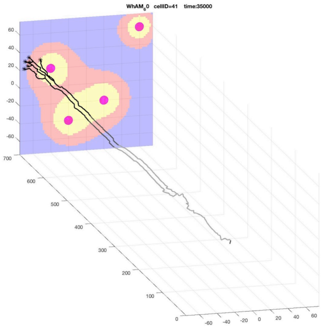

##### Lineage tree of survivors

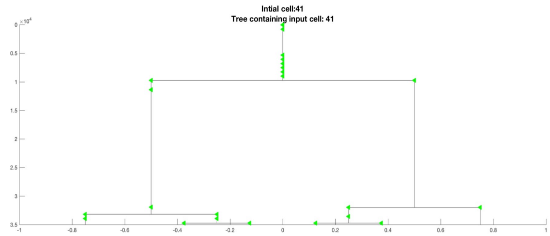


**Figure S16.** The full lineage tree for the initial cell #41 (top left), the lineage tree of survivors (bottom left), and the 3D spatio-temporal routes of all survived cells initiated from cell #41.

### Supplementary Figure S17

##### Cell # 43

##### Full lineage

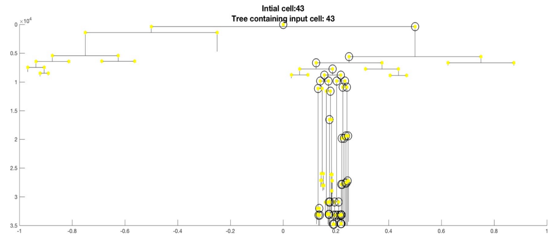


##### 3D lineage tree

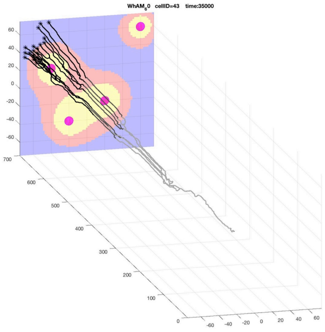

##### Lineage tree of survivors

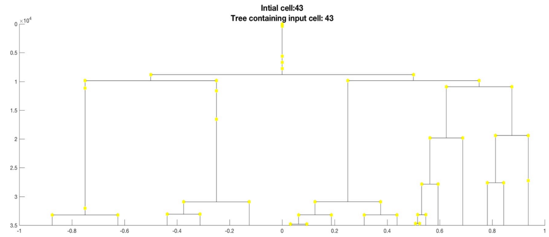


**Figure S17.** The full lineage tree for the initial cell #43 (top left), the lineage tree of survivors (bottom left), and the 3D spatio-temporal routes of all survived cells initiated from cell #43.

### Supplementary Figure S18

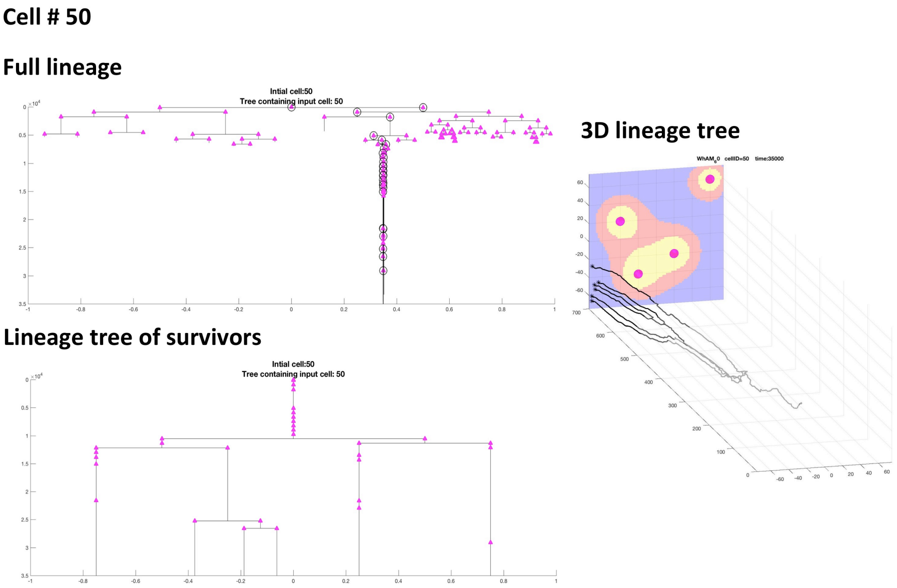


**Figure S18.** The full lineage tree for the initial cell #50 (top left), the lineage tree of survivors (bottom left), and the 3D spatio-temporal routes of all survived cells initiated from cell #50.
